# Tree-based automated machine learning to predict biogas production for anaerobic co-digestion of organic waste

**DOI:** 10.1101/2021.07.12.452124

**Authors:** Yan Wang, Tyler Huntington, Corinne D. Scown

## Abstract

The dynamics of microbial communities involved in anaerobic digestion of mixed organic waste are notoriously complex and difficult to model, yet successful operation of anaerobic digestion is critical to the goals of diverting high-moisture organic waste from landfills. Machine learning (ML) is ideally suited to capturing complex and nonlinear behavior that cannot be modeled mechanistically. This study uses 8 years of data collected from an industrial-scale anaerobic co-digestion (AcoD) operation at a municipal wastewater treatment plant in Oakland, California, combined with a powerful automated ML method, Tree-based Pipeline Optimization Tool, to develop an improved understanding of how different waste inputs and operating conditions impact biogas yield. The model inputs included daily input volumes of 31 waste streams and 5 operating parameters. Because different wastes are broken down at varying rates, the model explored a range of time lags ascribed to each waste input ranging from 0 to 30 days. The results suggest that the waste types (including rendering waste, lactose, poultry waste, and fats, oils, and greases) differ considerably in their impact on biogas yield on both a per-gallon basis and a mass of volatile solids basis, while operating parameters are not useful predictors in a carefully operated facility.

## INTRODUCTION

Anaerobic digestion (AD) has been used to generate combustible fuel from organic wastes since the 1800’s and, although advancements in synthetic biology have resulted in more targeted routes to producing specific fuel molecules, AD remains one of the most efficient strategies for converting mixed organic waste to renewable energy and nutrient-rich residual solids. In the U.S., there are over 2,200 biogas production sites, of which over 1,200 are industrial-scale AD facilities aligning with water resource recovery and additional 263 operate on livestock farms.^1–3^ Ambitious “zero waste” policies from local and state governments across the U.S. will require substantial new investments in infrastructure to divert organic waste from landfills, and AD is an essential part of any viable strategy.^4,5^ However, developing predictive modeling capabilities to estimate biogas yield and composition as a function of feedstock types has proved challenging. Unlike industrial bioreactors where a single microbial host utilizes a pure substrate (e.g. glucose), AD is most effective with heterogeneous organic feedstocks, as the use of a single feedstock (mono-digestion) often results a nutritional imbalance of substrates.^6–9^ Anaerobic co-digestion (AcoD) of multiple substrates has been widely employed to achieve the right nutrient balance and dilute inhibitory substances in the digester, improving biogas production and stability.^6–11^ Co-digestion can increase biogas yield by 25% to 400% compared to mono-digestion.^6,10^ AcoD is an effective practice for wastewater treatment plants (WWTPs), which receive organics for co-digestion with their sewage sludges in exchange for tipping fee revenue and combust the resulting biogas onsite to provide heat, electricity or mechanical power.^12,13^ Approximately 20% of AD facilities at U.S. WWTPs co-digest offsite wastes, with combined heat and electricity (CHP) as the dominant biogas use.^13^ The complexity associated with utilizing a microbial community grown on heterogeneous, variable substrates means that mechanistic modeling is usually impractical; the data required for a mechanistic model is vast and impossible to collect on a regular basis. Thus, biogas yield and composition prediction has remained a largely empirical exercise. Advanced regression techniques, under the umbrella of machine learning, offer an opportunity to improve upon current practices.

Generally, AD involves four successive steps that each rely on different communities of microbes, including hydrolysis, acidogenesis, acetogenesis, and methanogenesis.^7,14^ The resulting biogas product consists primarily of CH_4_ (50-75%) and CO_2_ (25-50%), with trace amounts of other components present as well such as H_2_O, O_2_, N_2_, H_2_S, NH_3_, siloxanes, and halogenated hydrocarbons.^15–18^ Biogas can be an attractive renewable fuel, particularly given the U.S. Environmental Protection Agency (USEPA) 2014 ruling that qualified biogas under the expanded Renewable Fuel Standard (RFS2) to claim D3 or D5 Renewable Identification Number (RIN) credits, depending on the type of organic wastes utilized.^19^ Biogas cleaning is necessary to remove trace contaminants, at which point it can be combusted for heat and/or electricity generation, used in gas fermentation processes, or upgraded and compressed or liquified for higher-value uses in transportation applications or steam methane reforming.^20,21^ Facilities employing AD to generate biogas must make decisions regarding tipping fees for, and willingness to accept, specific incoming waste types based on digester stability, biogas yield, and the likelihood of problematic contaminants (e.g. cutlery or other items that may clog or damage equipment). These decisions are often based on qualitative judgements and anecdotal observations rather than rigorous data analysis. Predictive modeling capabilities can provide a more quantitative basis to support decision-making for AcoD and improve resource utilization efficiency in the future.^6^

Mechanistic models, such as the well-known Anaerobic Digestion Model 1 (ADM1), have made important strides in the scientific community’s ability to predict digestion performance. However, the ADM1 model requires knowledge of many concentration state variables (i.e., the concentrations for detail components of substrates), which necessitates extensive ongoing analysis of substrates, thus limiting its applicability in industrial facilities where this data is not regularly collected.^22–24^ Also, the complicated microbial and physicochemical process of AD substantially affects the prediction accuracy of mechanistic models.^25^ Given the fact that AD is often a non-linear process, traditional statistical models (e.g., linear regression) have shown deficient performance for a generalized prediction of biogas production.^26^ When mechanistic modeling is not feasible or sufficient, and training data is available, machine learning (ML) can be the best option for developing predictive models and developing insights into the influence of key parameters. In the past decade, different ML techniques (summarized in Table S1) have been leveraged to predict biogas production, including connectivism learning (e.g., artificial neural network) and statistical learning (e.g., random forest, extreme gradient boosting, support vector machine).^22,23,27–37^ Previous research either employed a single technique or aimed to compare several techniques to select the best-performing approach (Table S1). These prior studies were also limited by the training data available to them.

Most ML studies have used fairly limited datasets, based on digesters operating at the lab scale or larger digesters fed with only a few substrates (Table S1). To the best of our knowledge, only the study from De Clercq et al. used a comparatively large dataset, with 4 years of operational data from an industrial AcoD facility treating 16 types of organic wastes.^23^ The team analyzed their data using elastic net, random forest, and extreme gradient boosting models to predict biomethane production.^23^ Our study aims to improve upon prior studies by using the most extensive dataset documented to-date, collected from an integrated full-scale WWTP-AcoD system spanning an 8-year operation period and accepting 31 different waste streams to produce biogas. By using a larger, more diverse dataset, the resulting model should provide greater insights and predictive capability, given ML’s usefulness for interpolation and the challenges in using ML models trained on limited datasets for extrapolation. This study also demonstrates a newer ML modeling approach – automated ML pipelines – that can be more easily replicated by practitioners and non-experts. One of the most successful automated ML systems is Tree-based Pipeline Optimization Tool (TPOT), which relies on genetic programming (GP) to recommend an optimized analysis pipeline including supervised classification/regression operators, feature preprocessing operators, and feature selection operators.^38–41^ We compared the performance of our TPOT model with more traditional ML techniques to understand how the results may vary and what insights about industrial AcoD operations can be gleaned from these approaches.

## METHODS

### Data collection and structure

Data used in this study were collected from East Bay Municipal Utility District’s WWTP plant (Oakland, California, U.S.) spanning an 8-year operation period (equivalent to 2,813 days). The EBMUD service area previously included multiple large industrial facilities that contributed to high biological oxygen demand (BOD) load at the WWTP, including a dog food factory and numerous canneries across Berkeley, Emeryville, and Oakland, California. The Resource Recovery program was developed to compensate for reduced BOD load as these industrial facilities shut down by accepting various high-strength non-hazardous organic trucked wastes to increase onsite energy production. A simplified process flow diagram for this integrated WWTP-AcoD system is shown in Figure 1. The liquid wastewater treatment processes mainly include coarse and fine screens, primary sedimentation tanks, aerated activated sludge basins, and clarifiers. Treated effluent is disinfected, dechlorinated, and discharged to the San Francisco Bay. The solid treatment processes include activated sludge thickeners, blend tanks (for solid blending of primary sludge, thickened waste activated sludge (TWAS), and trucked wastes), low thermophilic anaerobic digestion, and digested biosolids dewatering. Ferric chloride (FeCl_3_) is added to blend tanks (digester feed) for sulfide control and H_2_S reduction in the biogas. The biogas is combusted onsite to provide heat and electricity via a CHP system. The biosolids are used for land applications on non-edible crop sites or alternative daily cover at landfills.

**Figure 1.**
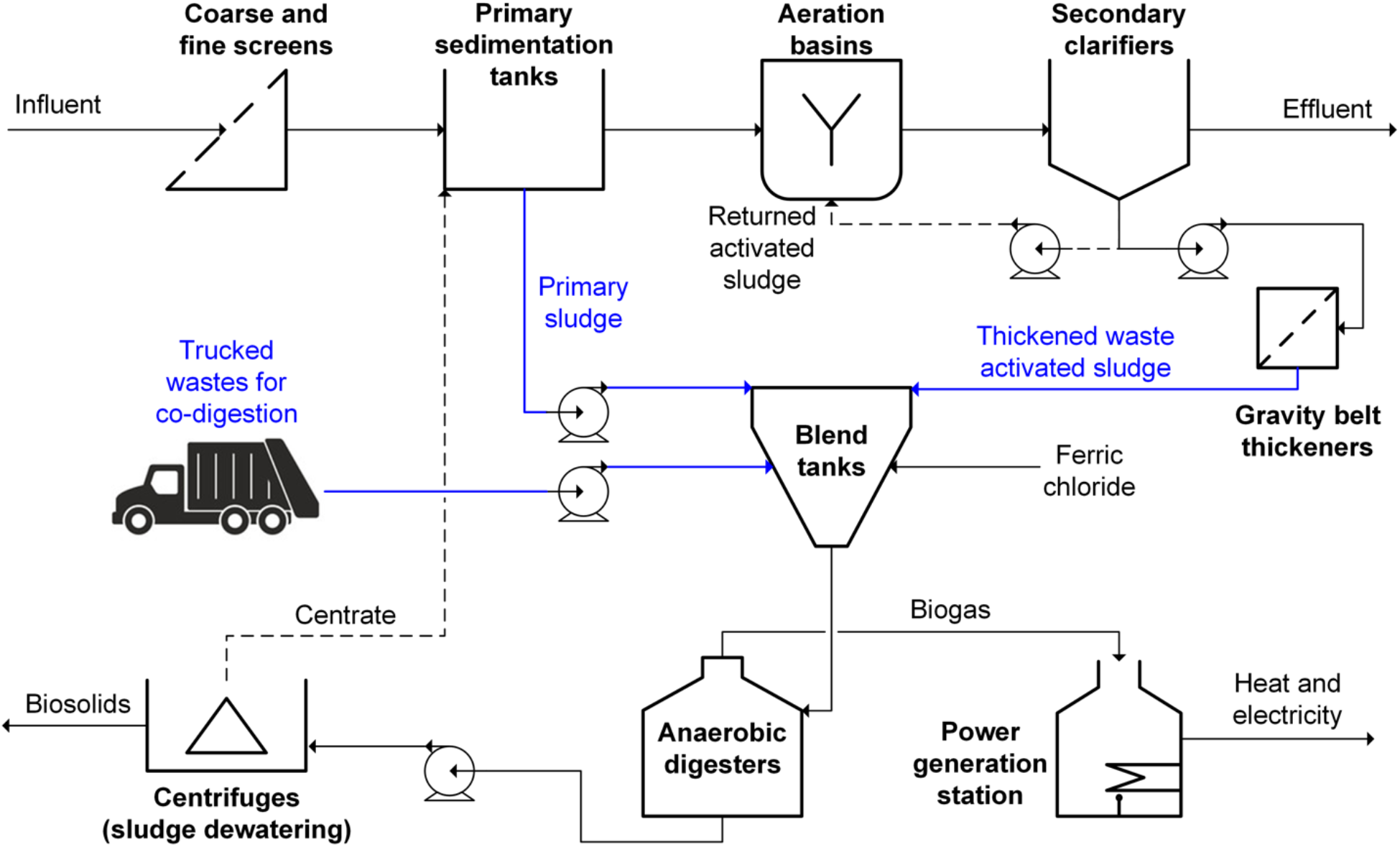
Simplified process flow diagram for the integrated full-scale WWTP-AcoD system. The waste streams fed into the AcoD facility are indicated in blue.

The AcoD facility accepts 29 types of trucked wastes, including waste categories such as brine, dairy, FOG (fats, oils, and greases), protein, process water, septage, sludge, food waste, winery, and general category for high-COD (chemical oxygen demand) organics (denoted as Organic(H)) and low-COD organics (Organic(L)). Two additional waste types are sourced from the WWT facility itself, for a total of 31 waste inputs. Definitions and brief descriptions of all waste streams fed into the AcoD facility is compiled in Table S2. Over the 8-year operation period, the daily input volume of each waste stream was recorded. 239 kilogallon/day of wastes on average were co-digested in the facility while the average total feed reached 668 kilogallon/day counting MMWT sludge wastes (primary sludge and TWAS). Also, the data for digester operating parameters were collected, including total solid (TS, %), volatile solid (VS, %), the content of volatile fatty acids (VFA, mg/L), and alkalinity (ALK, mg/L). TS and VS in the digester were calculated based on the daily input volume of each waste stream and its specific TS or VS level obtained from the composition analysis (Figure S1). Note that VFA and ALK were not measured every day, the missing values of VFA and ALK were imputed with the median value (raw data with non-normal distribution) and the mean value (raw data with normal distribution), respectively.

The model aims to predict the biogas production, thus the output (i.e., the target variable) is represented by biogas yield in standard cubic feet per minute (scfm). The input variables (i.e., features) to construct the dataset include the daily input volumes of 31 waste streams (primary sludge, TWAS, and 29 types of trucked wastes) and 5 operating parameters (TS, VS, VFA, ALK, and VFA/ALK ratio). Considering the digestion time, additional input variables were derived by lagging the daily input volumes of 31 waste streams for a period of time, namely, 0 (no lag), 1, 3, 5, 10, 20, and 30 days, respectively. In this way, a total of 222 input variables (31 waste streams × 7 time lags + 5 operating parameters) were created. This model configuration enables the following questions to be answered: which waste stream(s) have the greatest impact on biogas yield? On what timescale do these waste inputs impact the biogas yield (measured in days)? Figure 2 displays the raw data distribution for primary inputs and output. The model data structure is presented in Figure 3.

**Figure 2.**
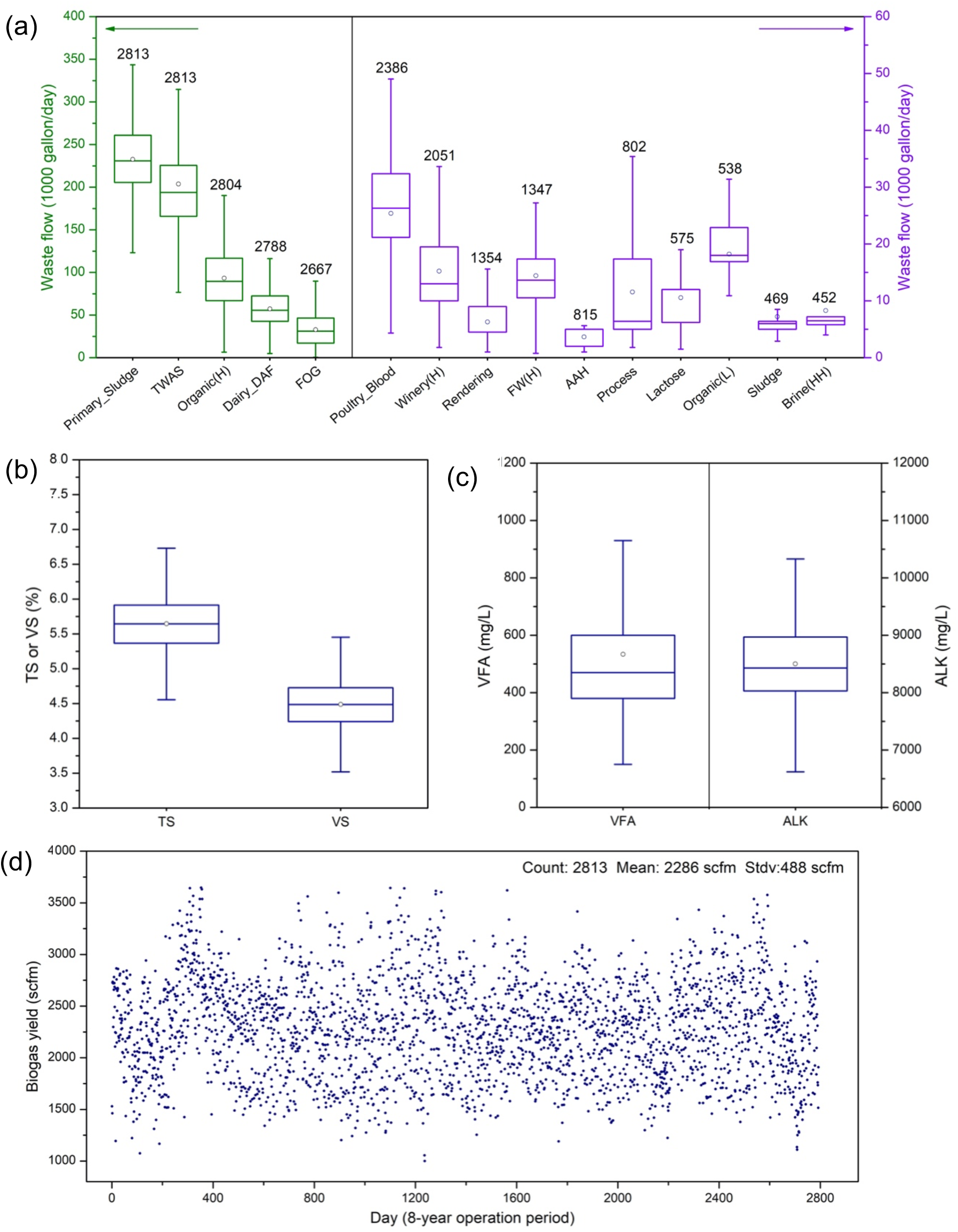
(a) Box plots (minimum, 25^th^ percentile, median, 75^th^ percentile, maximum, and mean by circles) displaying the data distribution for daily input volume (gallon/day) of 15 waste types that are in significant quantities. The flows for Primary_Sludge, TWAS, Organic(H), Dairy_DAF, and FOG are shown in the left y-axis (green) while those for other waste types are shown in the right y-axis (purple). The numeric values above the box refer to the number of collected data points, i.e., the number of days the digester accepts each waste type over an 8-year operation period. The name acronyms of wastes used along the x-axis are defined in Table S2. (b-c) Data distribution for digester operating parameters, including total solid (TS, %), volatile solid (VS, %), the content of volatile fatty acids (VFA, mg/L), and alkalinity (ALK, mg/L). (d) Evolution of the biogas production (scfm) during an 8-year operation period.

**Figure 3.**
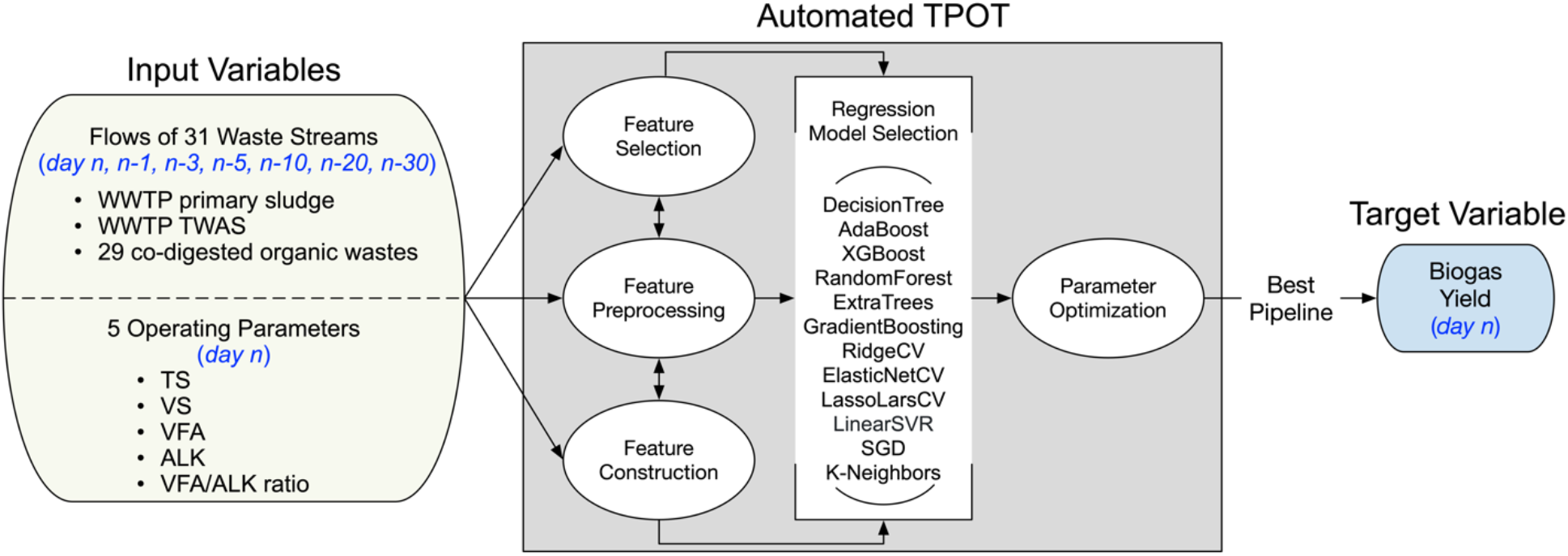
Schematic methodology using TPOT for biogas yield prediction based on 222 input variables, including 31 waste streams across different time lags (0, 1, 3, 5, 10, 20, 30 days) and 5 operating parameters at the current day. Typical steps in the ML pipeline automated by TPOT involve data transformation (feature selection, feature preprocessing, feature construction), model selection, and parameter optimization. The supervised learning regression models in the default TPOT configuration include decision trees, ensemble models (AdaBoost, XGBoost, forests of randomized trees, gradient tree boosting), cross-validated linear models (Ridge, Elastic Net, and Lasso using LARS algorithm), linear support vector regression (SVR), stochastic gradient descent (SGD), and k-nearest neighbors.

### TPOT overview

TPOT uses the dataset as input and recommends a best-performing ML pipeline with a series of operations related to feature selection, feature preprocessing, feature construction, and ML modeling (Figure 3). GP is used to optimize the pipelines. In this case, the population consists of a set of randomly generated pipelines to be evaluated; copies of the best-performing pipelines from each iteration (known as a generation) of the optimization process are created and imposed with random changes (e.g., the addition or removal of an operation or the parameter tuning of an operation), enabling the development of new pipelines that are never explored. The worst-performing pipelines are removed from the population at the end of each generation before starting the next generation. TPOT was run with default configuration and considered the following supervised learning regression models during the optimization process: decision trees, ensemble models (AdaBoost, XGBoost, forests of randomized trees, gradient tree boosting), cross-validated linear models (Ridge, Elastic Net, and Lasso using LARS algorithm), linear support vector regression (SVR), stochastic gradient descent (SGD), and k-nearest neighbors. Prior to the TPOT analysis, the dataset was randomly partitioned into a training set and a test set with a 0.75/0.25 train-test split ratio. The pipeline was trained on the training set and evaluated on the test set. The following TPOT parameter settings were used to generate a regression predictive model on the training set: the number of generations was 100, the size of population was 100, it used 5-fold cross validation (CV), and the scoring function (performance metric) was mean squared error (MSE). Further increasing in the number of generations and the size of the population did not improve the internal CV score. Further detail on the TPOT algorithm and its implementation can be found in previously published literature.^38–41^

### Model evaluation

The generalization performance of the trained model was evaluated on the test dataset. The metrics used to examine the precision and accuracy of the model include the coefficient of determination (R^2^) and the root mean square error (RMSE). While R^2^ offers a relative measure of model fit, RMSE provides an error metric in the same unit as the targe variable, making it highly interpretable. Higher R^2^ and lower RMSE of the predicted-versus-observed plots for the test dataset refer to higher precision and accuracy of a model for predicting biogas yield.

### Feature importance and partial dependence calculations

The two most widely used model interpretation techniques are feature importance and partial dependence (PD) plots. The permutation feature importance was calculated for the TPOT model using the scikit-learn library.^42^ Feature importance quantifies the relative importance of a feature (i.e., an input variable) by calculating the change in prediction error after the values of a given feature are randomly permuted. A larger increase in error suggests that the model relies on this feature to predict the target variable and thus has higher importance. MSE was used as the scoring function. The mean and standard deviation of feature importance were calculated over 50 permutations of a given feature in the training dataset. PD plots and individual conditional expectation (ICE) plots were produced using the PDPbox library.^43^ Using a previously fit model, PD plots visualize the predicted response as a function of the chosen feature while the effects of all other features in the model are averaged out. ICE plots show the functional relationship between the feature and the predicted response separately for each instance. Specifically, an ICE plot produces one line per instance, while the PD plot is simply the average of all lines in the ICE plot.

## RESULTS AND DISCUSSION

A critical first question to be answered through this analysis is whether machine learning can be effectively used to predict biogas yield based on detailed operating data. If the model achieved acceptable performance, a follow-on question is what insights the analysis provides into potential strategies for optimizing the facility studied here and other similar AD facilities.

### Prediction of biogas yield using TPOT regression model

To address the question of whether machine learning can be utilized to predict biogas yield, the TPOT model prediction performance was assessed by comparing the predicted values with the observed (measured) biogas yield in the test dataset (Figure 4). R^2^ and RMSE were used as the metrics to examine the precision and accuracy of the model, respectively. The regression model selected in the best-performing TPOT pipeline was extremely randomized trees (ExtraTrees). The ExtraTrees pipeline from TPOT performed well on the test dataset with an R^2^ of 0.72, which generally represents good predictive capacity for a model trained on real-world industrial data.^25^ For the sake of comparison, an alternative model using artificial neural network (ANN) was also developed. ANN has been the most commonly used technique to predict the performance of AD facilities (Table S1). In this case, the most popular ANN type, multi-layer perceptron (MLP), was employed (Figure S2) as the baseline for comparison with the TPOT model. TPOT (R^2^ = 0.72, RMSE = 247 scfm) outperforms MLP (R^2^ = 0.56, RMSE = 327 scfm). Notably, while many ANN parameters require manual tuning, TPOT automates the parameter tuning process, making it a more practical ML approach for non-experts.

**Figure 4.**
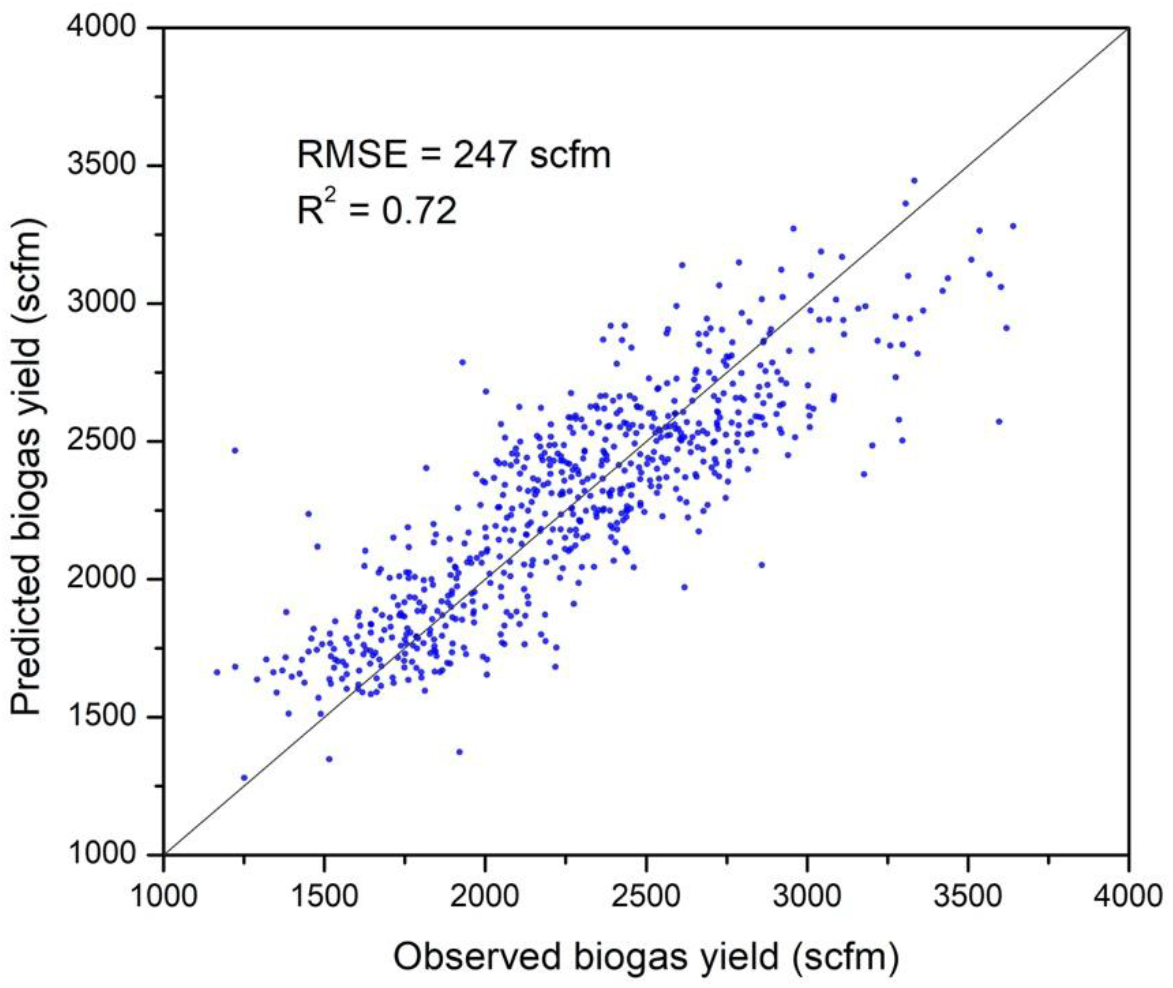
Comparison between observed and predicted biogas yield (scfm) for the test dataset using TPOT. ExtraTrees regression model was selected for the best prediction performance. Model prediction performance is evaluated by RMSE and R^2^, and also visually revealed by the extent of data clustering around the identity line (y = x, in black).

### The importance of input variables influencing biogas yield

Model interpretability is critical, whether it is developed as a research tool or to guide decision-making for facility operation. This study relies on expertise in bioprocesses and experience in the operation of AD facilities to guide the model development and interpretation such that the results answer scientifically interesting and operationally relevant questions. To investigate the influence of input variables (daily input volumes of 31 waste streams each evaluated over different time lags and 5 operating parameters) on biogas yield, permutation feature importance was generated for the TPOT model (Figure 5). The importance scores provide insights into which input variables have the greatest influence on biogas yield.

**Figure 5.**
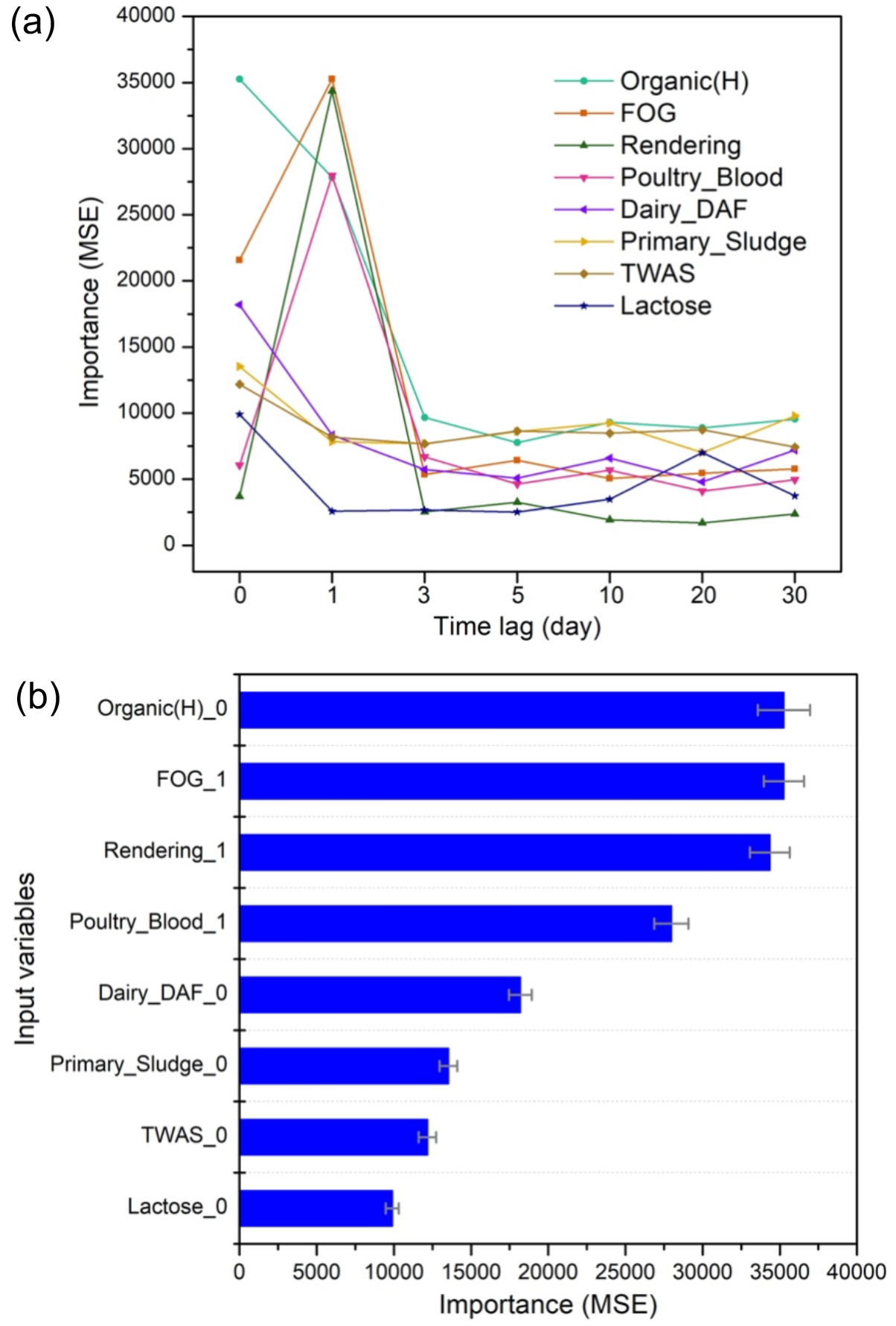
(a) Input variable importance of top 8 waste types in the TPOT model across different time lags (0, 1, 3, 5, 10, 20, 30 days). The importance values of all input variables are compiled in Table S3. The relative importance scores are calculated using the permutation feature importance technique. An input variable that leads to a higher increase in the mode prediction error (mean squared error, MSE) upon randomly permuted is more important in the model. The name acronyms of wastes are defined in Table S2. (b) Importance ranking for the 8 most influential waste types, of which each shows the highest importance score from the time-lag variable set (data compiled from (a)). The suffix on waste names represents the specific number of lagged days. Error bars denote the standard deviations over 50 times of permuting an input variable.

Operating parameters, while not truly independent of other input variables like feedstock inputs, can improve the model performance. The 5 operating parameters that are routinely monitored at the EBMUD facility studied here include TS, VS, VFA, ALK, and VFA/ALK ratio, and all are essential factors for digester design considerations and process stability. TS, representing the percentage of the dry matter (organic or inorganic), is an important attribute of digester design and operation. For example, higher TS usually results in smaller digester size and lower heating demand.^44^ VS is typically regarded as a measurement of organics, serving as the basis for determining the digester organic loading rate. VFA is generated from the acidogenesis stage, comprising a class of organic acids (e.g., acetic acid, propionic acid, butyric acid). Neutral pH is optimal for methanogens.^14^ VFA acidification is widely considered to be a cause of digester failure because methanogens are sensitive to low pH, which has an inhibitory effect on their growth.^14,44^ As a result, ALK is needed to provide buffer capacity to neutralize VFA, thus controlling pH. A balance between VFA production and consumption by ALK (reflected by VFA/ALK ratio) is critical for a stable AD process.

The most important input variables that affect biogas yield are waste inputs from high-COD organics, FOG, dairy, protein, lactose, and sludge categories, while operating parameters did not have high importance scores (Figure 5, Table S3, also see Table S2 for the waste categories and the definitions of waste name acronyms). The less significant role of operating parameters in influencing biogas yield can be attributed to the fact that the EBMUD facility is carefully managed such that conditions do not frequently deviate from the acceptable range, therefore, biogas production is predominantly regulated by the types of waste being fed into the digester. At a facility in which pH, for example, is allowed to become sufficiently acidic to inhibit microbial growth, the resulting model might show a stronger relationship between VFA and biogas yield. The top 8 waste types include Organic(H), FOG, Rendering, Poultry_Blood, Dariy_DAF, Primary_Sludge, TWAS, and Lactose. The appropriate time lag to assign for each waste type was determined, representing the duration between when it is delivered to the facility and when it has the greatest impact on biogas yield. Figure 5a shows the importance results for each waste type and time lag. Poultry blood, rendering waste, and FOG all appear to have the greatest impact on biogas yield with a one-day time lag, whereas all other waste types have the largest importance scores with no time lag (zero days). This agrees with the facility operators’ observations that the biogas yield is obviously boosted within a matter of hours after feeding the more sugar-rich wastes into the digesters while blood and other protein and lipid-rich wastes typically boost yields within a day.

Figure 5b compiles the best-performing version of each input variable (based on the time lag analysis in Figure 5a) and their relative performance was compared to determine which waste types are the primary drivers of biogas yield. Organic(H), while not the largest input by volume (Figure 2a), results in the highest importance score. This is a general category used by the facility to denote waste streams with a COD greater than 20,000 mg/L. Organic(H) might include, for example, carbohydrate-rich waste from beverage processing facilities. Anecdotally, sugar-rich wastes have been noted to produce a near-immediately evident impact on microbial activity in the digesters. Followed by Organic(H) in terms of impact on biogas yield are the three protein and lipid-rich waste types, all of which are well known to be desirable supplemental inputs in wet AD.^45,46^ In addition, the results indicate that, although WWTP sludge wastes (primary sludge and TWAS) have considerably higher daily input volumes than the trucked co-digested wastes (Figure 2a), their contributions to biogas yield are less significant than the wastes in the categories of high-COD organics, FOG, dairy, and protein. Compositional analysis on the co-digested wastes shows that the wastes in these categories typically have higher TS, VS, COD, and total nitrogen than others (Figure S1). Combined, the waste types with higher organic matter contents and faster digestion rates are more contributory to biogas yield.

### The quantitative relationships between the most influential waste inputs and biogas yield

While feature importance calculations identify the most influential input variables, it is necessary to explore the functional relationships between these important input variables and the target variable. PD (Figure 6) or ICE (Figure S3) plots enable us to isolate the quantitative effect of adjusting the values of an input variable on future prediction outcome. PD plots visualize the marginal (average) effect of an input variable of interest on the target variable while ICE plots demonstrate the heterogeneity or dispersion of the effect.^47^ The relative impacts of the most influential waste inputs (in daily volume) on biogas yield are illustrated by the PD plots (Figure 6a). As expected, all these waste inputs positively impact biogas yield. The slopes of PD lines reveal the increase in biogas yield on a unit basis of each waste input throughout its whole value range, following the order: Rendering > Lactose > Poultry_Blood > FOG > Organic(H) > Dairy_DAF > Primary Sludge > TWAS. The results between feature importance and PD calculations are consistent, considering that the overall effect of each waste on biogas yield is ascribed to its specific biogas production capacity (revealed by the PD line slope) as well as its feed availability (the feed amount). High-COD organics (Organic(H)) emerge as having the highest importance score, but after examining per-unit-flow values, rendering waste, lactose, poultry blood, and FOG all result in greater increases in biogas yield.

**Figure 6.**
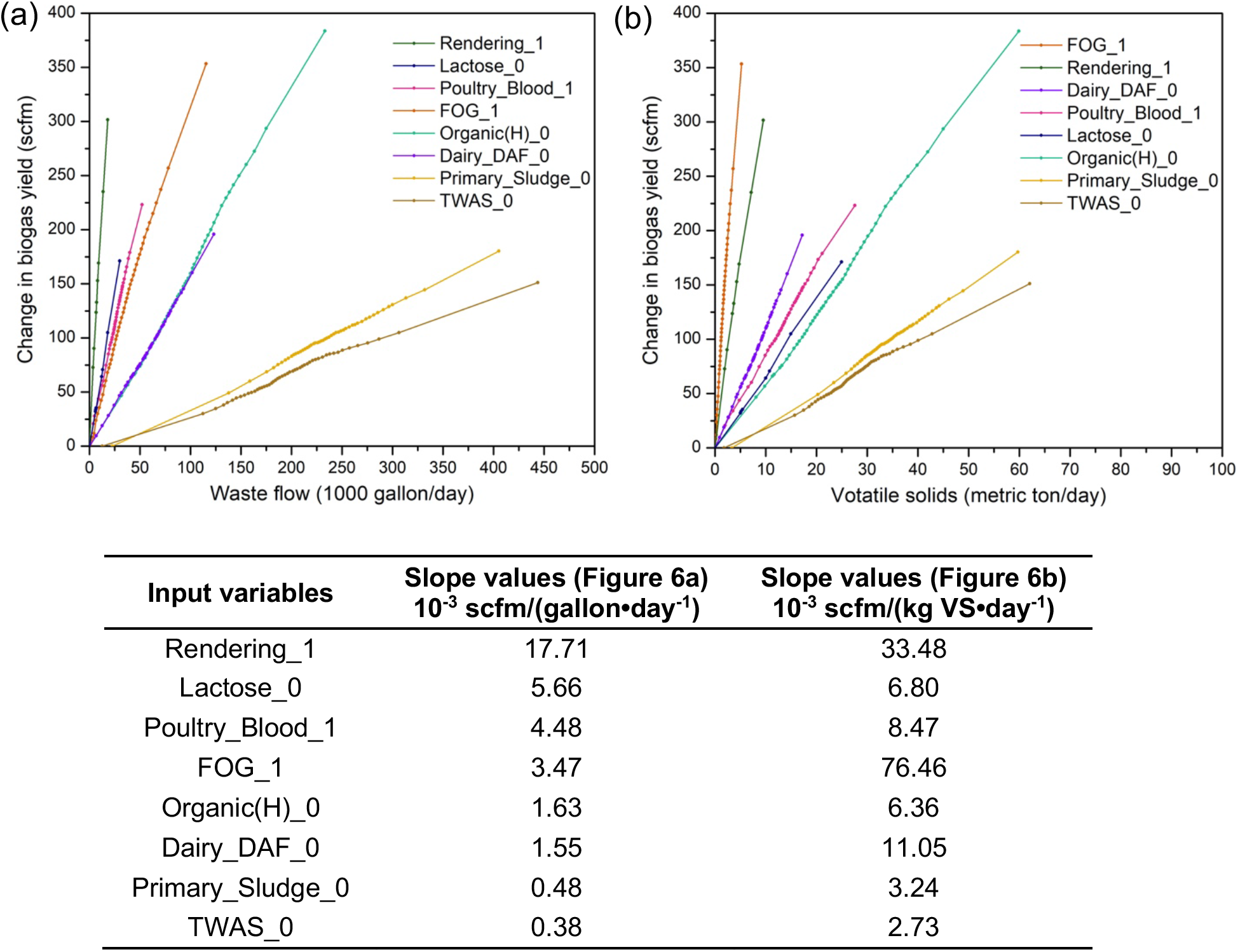
Partial dependence (PD) plots depicting the quantitative relationships between (a) the daily input volume (gallon/day) or (b) the daily volatile solids (VS) load (metric ton/day) of the 8 most influential waste types and the resulting change in biogas yield (scfm). The table provides the numeric values of line slopes determined by linear curve fitting. The legends are arranged in the descending order based on the approximated slopes of the lines. Each line is for the input variable that shows the highest importance score in the time-lag set (0, 1, 3, 5, 10, 20, 30 days) of each waste type. The suffix on waste names denotes the specific number of lagged days. The dots on the lines refer to 50 percentile points across the whole value range of each input variable, note that some percentile points can be same thus each line does not necessarily contain 50 points. The dispersion of dots visually reveals the data points distribution of each input variable. Individual conditional expectation (ICE) plots for these 8 input variables are shown in Figure S3.

Because different waste streams contain varying amounts of solids, AD facility operators often characterize wastes on the basis of VS. Normalizing the results in Figure 6a with respect to the daily VS load provides the results in a more usable (and generalizable) form for other facilities (see Figure 6b). The normalized results in Figure 6a suggest considerable variation in the impact on biogas yield on a mass of VS basis. The ordering, from greatest impact to least impact on biogas yield also shifts; FOG, rendering waste, dairy waste after dissolved air flotation (DAF) treatment, poultry blood, and lactose all demonstrate higher capacity for biogas production on a per-mass VS basis. Together, the feature importance ranking and PD plots provide useful insights into how different inputs impact biogas yield that can be leveraged for plant operators to manage and operate the AcoD facility. The numeric slopes in Figure 6 can be used to predict the impact of any given waste stream on biogas yield, and thus inform tipping fees and priorities for which wastes to accept. The fact that some wastes impact biogas production more rapidly than others indicates that operators could attempt to manage incoming waste such that biogas production is as stable as possible, thus minimizing the need for flaring excess biogas. However, anecdotal evidence from the EBMUD facility and other local AcoD and AD facilities suggests that the greatest challenge in timing is the reduction in waste hauling during weekends. Facilities could make use of a combination of on-site liquid storage and slower-degrading waste types to partially mitigate this problem.

### The opportunity to predict biogas trace compounds using ML techniques

Raw biogas typically contains trace undesirable compounds (contaminants), of which the amounts are highly dependent on the origin of organic sources. H_2_O, H_2_S, NH_3_ are commonly present while siloxanes and halogenated hydrocarbons can also be present.^15–18^ These contaminants can cause problems to the equipment (e.g., engines, pipelines, valve fittings) for biogas utilization. For example, H_2_O, H_2_S, NH_3_, and halogenated hydrocarbons cause corrosion SiO_2_ formed from siloxanes causes abrading gas motor surfaces.^15,18,48^ Biogas cleaning is thus normally conducted for all commonly used biogas applications. The prediction of these trace compounds, especially by employing ML techniques, can be informative if it enables operators to select waste streams that minimize their formation or simply optimize gas cleaning investments to manage the contaminants.

H_2_S is the most influential trace compound to be treated in biogas for energy applications, considering the risk of sulfide stress cracking (embrittlement)^49^ and the ubiquitous presence of sulfur in biological substrates (particularly those with high protein levels).^17^ The methods for biogas cleaning differ according to the required quality demands for the contaminants in specific end uses of biogas. H_2_S is considered as the main component for assessing the biogas quality in heating boilers and internal combustion engines, where its concentration should be lower than 1,000 ppm.^17,49^ However, for biogas use as vehicle fuel or natural gas, the requirements for H_2_S become much stricter (e.g., < 120 ppm in the U.S.) and maximum allowable concentrations vary by country.^17,49^ H_2_S is typically removed during digestion through a precipitation reaction with metal ions (Fe^2+^, Fe^3+^) or after digestion by absorption (e.g., using water for scrubbing or metal oxide), adsorption (e.g., on activated carbon), biological filters, and membrane separation.^15,16^

The EBMUD facility studied here removes H_2_S by adding FeCl_3_ to the digester, which reduces the H_2_S concentration to less than 300 ppm and thus satisfies the requirement for combustion on site. This also means that H_2_S concentrations are measured in the biogas after much of the sulfur has been removed by FeCl_3_. Furthermore, the facility monitors H_2_S on an approximately weekly basis, so the dataset for H_2_S is fairly limited in its granularity. As one might expect, using ML techniques to predict H_2_S did not prove to be useful because the H_2_S output is most strongly correlated with the dose of FeCl_3_. Since the actual H_2_S content produced from the input wastes is not measured, the minimum required amount of FeCl_3_ cannot be accurately estimated with our modeling approach. That said, ML techniques do have the potential to predict concentrations of trace compounds in such a way that could be useful in applications requiring high biogas quality, where frequent measurement of biogas composition is conducted. First, the in-situ H_2_S removal methods during digestion are less efficient in achieving the required level of H_2_S (and other contaminants of concern) for transport fuel or pipeline quality, where post-treatment methods after digestion are needed.^16^ Also, very little work has been done to explore the potential of ML techniques for predicting these biogas contaminants. To our knowledge, only Strik et al. applied ANN (MATLAB Neural Network Toolbox) to predict H_2_S and NH_3_ concentrations in an experimental AD setup a decade ago.^50^ This topic presents an opportunity to use computational methods to improve the efficiency of the biogas industry and gain insights into the complex dynamics of the microbial communities that break down mixed organic waste. With richer datasets from facility operations and even lab-scale experiments, researchers could harness (automated) ML techniques to develop better predictions of the concentrations of trace compounds and biogas quality for biogas utilization systems that employ the post-treatment biogas cleaning methods (e.g., for biogas upgrading technologies). This could also help with the management of air pollutant emissions, as NOx emissions from flaring has been shown to have a relationship with NH_3_ concentrations in raw biogas.^51^ Tracking NH_3_ could also serve as a proxy for nutrient loading in the effluent, which could enable facilities to develop strategies that minimize water quality impacts of using high-nitrogen wastes.

Anaerobic digesters have an important role to play in diverting organic waste from landfills and producing renewable energy.^5^ AcoD technology in particular is being embraced as a pragmatic strategy for increasing biogas production, leveraging existing infrastructure, while overcoming the challenges associated with the substrate properties and system stability in single-substrate AD process. The key to the success of AcoD processes is system optimization, and an ability to manage a diverse set of incoming waste streams. ML models, which are ideally suited to capturing the behavior of systems that are too complex to model mechanistically, can improve researchers’ and operators’ understanding of the AcoD process and its performance as a function of varying feed substrates or operating conditions. Our work contributes to a growing field of biogas production prediction using ML techniques and the use of TPOT with a substantially larger dataset than any previously documented (based on 8-year industrial-scale operations) makes this study unique. Our work provides evidence for the robust predictive power of TPOT applied to AcoD modeling, as demonstrated by its superior prediction performance compared to the basic ANN model (MLP). The combination of feature importance and PD analyses allowed us to differentiate between waste streams that have a large impact because of greater incoming volumes and waste streams that have a greater impact per unit of waste input to the digester. Our approach of testing different time lags also provided insights into how different wastes are broken down once loaded into the digester. By developing and improving predictive models for AcoD performance, we hope to enable more efficient facility operation, better understanding of how microbial communities respond to different substrates and operating conditions, and ultimately a more sustainable organic waste valorization industry.

## ASSOCIATED CONTENT

### Supporting Information

The Supporting Information is available free of charge on the ACS publications website, including literature summary of ML applications in predicting biogas production from anaerobic digestion (Table S1), brief description on the waste streams fed into the AcoD facility (Table S2), typical composition analysis data on the waste categories (Figure S1), biogas yield prediction using ANN-MLP (Figure S2), ranked permutation importance values for all input variables (Table S3), and ICE plots for the 8 most influential waste types (Figure S3).

### AUTHOR INFORMATION

**Corresponding Author**

*

### Notes

The authors declare no competing financial interest.

## ACKNOWLEDGEMENTS

The authors would like to acknowledge Michael Hyatt and John Hake from the East Bay Municipal Utility District for providing access to their facility data and giving extensive feedback on the design and results of our study. This study was supported by the U.S. Department of Energy, Energy Efficiency and Renewable Energy, Bioenergy Technologies Office. This work was also enabled by tools and resources provided by the DOE Joint BioEnergy Institute (http://www.jbei.org) supported by the U.S. Department of Energy, Office of Science, Office of Biological and Environmental Research, through contract DE-AC02-05CH11231 between Lawrence Berkeley National Laboratory and the U.S. Department of Energy.

---

**FOR TABLE OF CONTENTS GRAPHIC ONLY:**

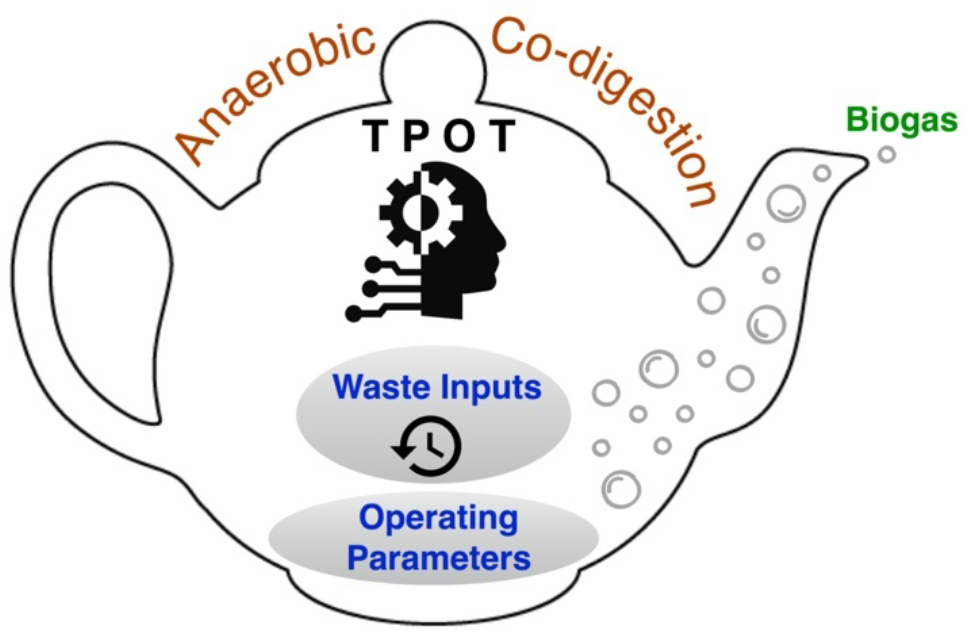

## SYNOPISIS

Leveraging automated machine learning techniques to predict biogas production can improve resource utilization and management efficiency advancing organic waste-to-energy industries.

## Notes

### Competing Interest Statement

The authors have declared no competing interest.

